# LocusZoom.js: Interactive and embeddable visualization of genetic association study results

**DOI:** 10.1101/2021.01.01.423803

**Authors:** Andrew P. Boughton, Ryan P. Welch, Matthew Flickinger, Peter VandeHaar, Daniel Taliun, Gonçalo R. Abecasis, Michael Boehnke

**Affiliations:** Department of Biostatistics and the Center for Statistical Genetics, University of Michigan, Ann Arbor, MI; Regeneron Pharmaceuticals, Tarrytown, NY

## Abstract

LocusZoom.js is a JavaScript library for creating interactive web-based visualizations of genetic association study results. It can display one or more traits in the context of relevant biological data (such as gene models and other genomic annotation), and allows interactive refinement of analysis models (by selecting linkage disequilibrium reference panels, identifying sets of likely causal variants, or comparisons to the GWAS catalog). It can be embedded in web pages to enable data sharing and exploration. Views can be customized and extended to display other data types such as phenome-wide association study (PheWAS) results, chromatin co-accessibility, or eQTL measurements. A new web upload service harmonizes datasets, adds annotations, and makes it easy to explore user-provided result sets.

**Availability:** LocusZoom.js is open-source software under a permissive MIT license. Code and documentation are available at: https://github.com/statgen/locuszoom/. Installable packages are also distributed via NPM. Additional features are provided as standalone libraries to promote reuse. Use with your own GWAS results at https://my.locuszoom.org/.

## Introduction

Genetic association studies are a workhorse technique in the study of complex disease genetics, and in recent years the numbers of reported studies and associations have risen dramatically. (Buniello et al., 2019) It has long been recognized that interpreting genetic association studies requires significant context, including linkage disequilibrium (LD) patterns, recombination rate, nearby genes, results for related traits, chromatin accessibility, and other information. LocusZoom (Pruim et al., 2010) is now the standard tool for visualizing GWAS results in the context of relevant genetic annotations, with both command line and web-based versions. Here, we describe a new JavaScript implementation of LocusZoom that significantly expands its capabilities, supporting interactive web-based visualizations with options for both public and private datasets.

## Key features

The basic LocusZoom plot has been described elsewhere. (Pruim *et al.*, 2010) Briefly, the main area presents a scatter plot of association p-values vs chromosomal position, frequently using color to summarize LD with the lead genetic variant. Additional tracks show recombination rate, nearby genes, and relevant annotations such as variants for which associations have been reported in the EBI GWAS catalog (Buniello *et al.*, 2019). The region size is user selectable, typically < 1Mb. (Figure 1)

**Figure 1:**
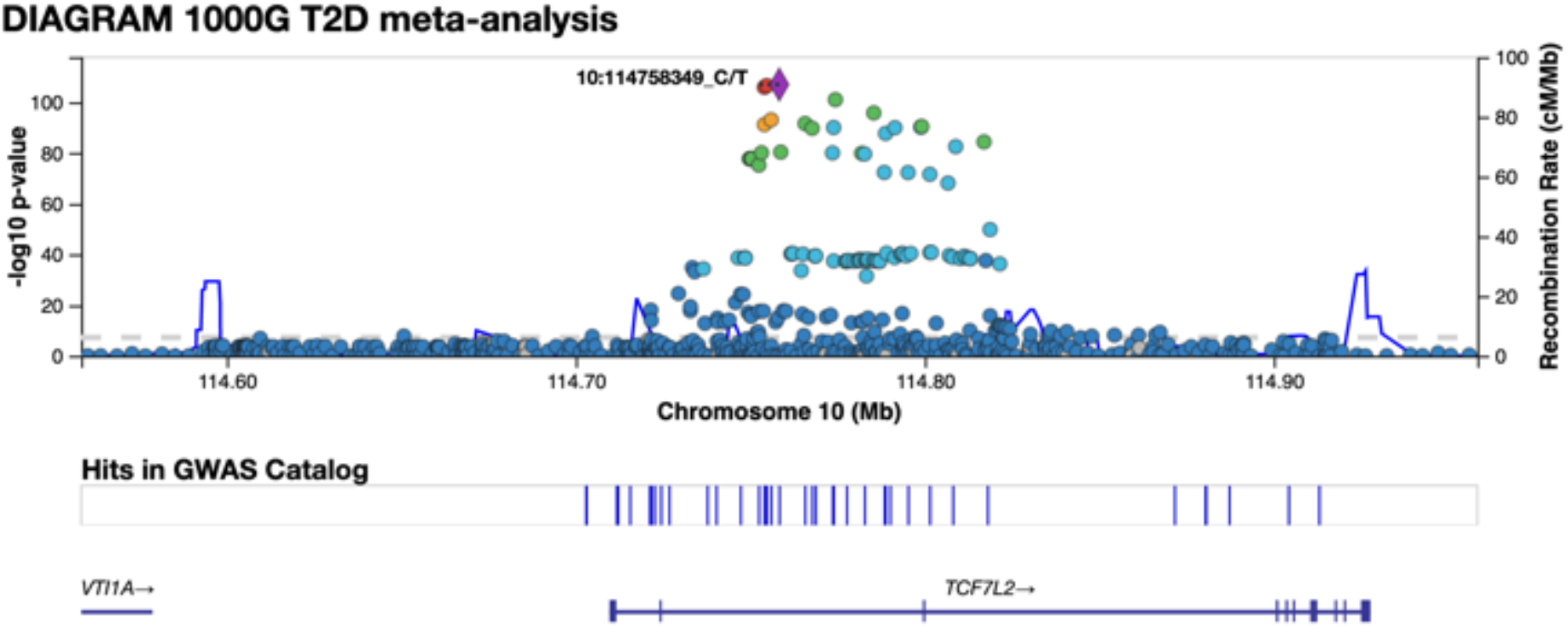
A standard LocusZoom region plot. Association summary statistics are presented as a scatter plot, with point colors indicating LD relative to the lead (most significant) variant in the region. Tick marks represent variants with significant hits in the GWAS catalog. Individual points can be labeled or annotated interactively via tooltips that expose additional information. Genes indicate introns and exons (horizontal and vertical bars)

LocusZoom.js provides an embeddable JavaScript widget, with all data rendering handled client side using web-standard technologies such as HTML, CSS, and SVG. This allows significant dynamic behavior. For example, the user may hide certain elements (e.g. pseudogenes), recolor the plot based on a different LD reference population, or toggle to visualize a credible set. To facilitate exploration of a genome-wide dataset with many significant loci, the region in view may be changed by user gestures (drag and zoom) or programmatically. At any time, the current view may be exported as PNG or high-resolution SVG files. Event callbacks allow the plot to trigger and respond to external events, such as updating a data table to reflect the information shown on the plot. Within the plot, all data elements provide tooltips that expose additional information or functionality.

LocusZoom.js most commonly fetches data via REST APIs, and we provide an API server (https://portaldev.sph.umich.edu/docs/api/v1/) with access to standard datasets such as 1000 Genomes Project LD, GENCODE genes, HapMap-based recombination rate, and the EBI GWAS Catalog. All annotations are provided for genome builds GRCh37 and GRCh38. A plugin-based architecture makes it straightforward to add data adapters for other formats, including local and remote tabixed files. In practice, this allows plots to be generated from a static file host such as S3, reducing the barrier to entry. Since rendering is performed in the browser, the only dependency is the embedded code; private datasets do not need to be transmitted to a third-party server.

When LocusZoom.js is integrated into existing web applications, significant customization is possible due to a track and plugin-based architecture, with a declarative layout system that can be expanded to support additional data types. LocusZoom.js has already been used to visualize PheWAS, eQTL, chromatin co-accessibility, and chromatin state annotations; an example gallery demonstrates many visual and interactive features (https://statgen.github.io/locuszoom/). Track types can be layered, stacked, and dynamically added to foster exploration. LocusZoom.js has been incorporated into numerous third-party sites including the Type 2 Diabetes Knowledge Portal (https://t2d.hugeamp.org/) and PheWeb (Gagliano Taliun *et al.*, 2020). (see Supplementary information for further examples)

## Interactive plots of user-provided data

Although originally developed for use in centralized data sharing portals, LocusZoom.js can also be used by researchers with their own datasets. We have created a web-based tool called LocalZoom (https://statgen.github.io/localzoom/) that allows interactive plotting of user-provided datasets from a local, tabix-indexed (Li, 2011) file. This occurs in the web browser and no data are uploaded to a central server, thus protecting privacy of sensitive results. Upon selecting the file, a modal dialog prompts the user to identify key fields such as chromosome, position, reference and alternate alleles, (−log_10_) p-value, effect size, standard error, and alternate allele frequency. Built-in heuristics and data validation allow LocalZoom to parse most user-provided files automatically. To aid exploration of results, LocalZoom exposes a number of advanced features (Table S1).

A search box allows easy access to genetic regions based on chromosome:position, gene name, or rsID syntax. A 95% credible set is calculated for the region in view based on a simple Bayesian refinement method (The Wellcome Trust Case Control Consortium *et al.*, 2012), and the visualization can be updated to reflect this calculation. Additional GWAS datasets can be added to the plot as vertically stacked panels to allow comparisons between studies, and clicking on a variant of interest will display associations from other phenotypes in a PheWAS of the UK BioBank (Zhou *et al.*, 2018). Data may be viewed and exported via an accompanying tabular view that includes dynamically calculated results such as credible set membership.

## Upload service with additional features

Due to the size and complexity of GWAS summary statistics, some additional features require an asynchronous pre-processing step outside the web browser. We have created a new, fully interactive upload service (https://my.locuszoom.org) that allows both public and private sharing, while incorporating all of the advanced display features from LocalZoom (Table S1). By allowing users to upload their files in advance, additional processing and annotations are possible, such as Manhattan and QQ plots, nearest gene and rsID labels, and a convenient batch view mode to browse the most significant hits across the entire genome. At present, uploaded files are limited to ~1GB in size. To protect sensitive data, a Google account is required to upload a study, but shared datasets can be viewed anonymously. A range of studies are provided for public browsing. In its first year of operation, this service has generated >102,000 plots across >7,700 datasets. In the future, we plan to support additional features such as custom user-defined annotations and comparisons to published studies.

## Acknowledgements

We are grateful to Chris Clark, Ben Alexander, Sarah A. Gagliano Taliun, and Parul Kudtarkar for many helpful discussions.

## Funding

This work was supported by grants HG009976 from the NIH and BOEH15AMP from the Foundation for the NIH.

## Conflict of Interest

G.R.A. is an employee of Regeneron Pharmaceuticals; he owns stock and stock options for Regeneron Pharmaceuticals.

