## Supplementary Information for "LocusZoom.js: Interactive and embeddable visualization of genetic association study results"

### LocusZoom.js: Usages and Examples

LocusZoom.js has been used for the display of many scientifically interesting datasets across a range of studies and organizations. It can be difficult to convey the interactive and exploratory data analysis capabilities of a tool through prose; the provided URLs offer examples of how LocusZoom.js is used on various sites across the internet, along with a description that highlights specific features of interest.

#### LocusZoom.js Example Gallery (https://statgen.github.io/locuszoom/)

- Demonstrates a range of features, including the capability to perform region-based aggregation tests (using raremetal.js) in the browser, using data from the METSIM study (Laakso *et al.*, 2017). The user is able to generate, explore, and interpret new hypotheses directly in the web browser; they are not limited to displaying pre-generated results.

#### Human Genetics Amplifier (HuGeAMP) / Knowledge Portal Network (https://t2d.hugeamp.org/)

- A family of interconnected data-sharing web portals, including the Type 2 Diabetes Knowledge Portal. Contains association results for hundreds of datasets and traits. Users can explore their own comparisons by adding custom tracks (such as accessible chromatin state annotations) to place results in additional biological context.
- A 2020 webinar provides an overview of key LocusZoom.js features in the T2D Knowledge Portal, including interactive calculation workflows: https://youtu.be/AmU7IIAeZJ4.

#### PheWeb: A tool for building multi-phenotype GWAS browsers (https://pheweb.org)

- Uses LocusZoom.js to render Phenome-Wide Association Study (PheWAS) and Genome-Wide Association Study (GWAS) visualizations. Users can interactively click any point on the scatter plot to move between different views of the dataset, and an additional track allows comparison of significant results to existing claims from the EBI GWAS catalog.
- PheWeb (Gagliano Taliun et al., 2020) is used as the basis of several other genome browsers, including the Oxford Brain Imaging Genetics (BIG) server (http://big.stats.ox.ac.uk) and FinnGen results browser (https://results.finngen.fi/about).

#### LocusZoom.org upload service (https://my.locuszoom.org/)

- Allows users to upload their own GWAS results and receive a shareable link to the exact view and region of interest.
- A credible set is automatically calculated in the web browser, demonstrating the ability to enhance datasets with new scientific results without requiring re-ingest of previously uploaded files.
- An interactive table allows the data associated with the plot to be exported for followup analysis; this table is synchronized with user interactions in the plot.

#### FIVEx eQTL Browser (https://eqtl.pheweb.org/)

- Explore GTEx eQTL results for single-variant and region views. Interactive controls allow the user to add or overlay annotations to better understand the origin of interesting signals.

### Implementation and Architecture

LocusZoom.js achieves significant flexibility by defining reusable display elements (data layers) that can be combined to meet the needs of the analyst. Each layer describes the basic data structures and rendering options required to produce a desired effect. Layers can be superimposed (such as a scatter plot and line of genome-wide significance) or aligned (panels) within a single connected plot area (Figure S1); changes in the viewable region will then propagate to all panels within that plot.

To ensure that elements are reusable, a separation of concerns is enforced between data adapters (data) and data layers (presentation). Custom features can be added via a plugin system; we provide several sample plugins that demonstrate usage of LocusZoom.js with custom data retrieval, specialized D3.js rendering types, etc. (Table S1) Support is provided for environments either with or without advanced JavaScript tooling available. Efforts are made to keep dependencies to a minimum.

A public API server (https://portaldev.sph.umich.edu/docs/api/v1/) provides access to several standard datasets: 1000 Genomes Project (The 1000 Genomes Project Consortium, 2015) LD, GENCODE genes (Frankish *et al.*, 2019), HapMap-based recombination rate (The International HapMap Consortium, 2003), and the EBI GWAS Catalog (Buniello *et al.*, 2019).

For the most up-to-date details of usage, please consult the documentation and example gallery at: https://github.com/statgen/locuszoom.

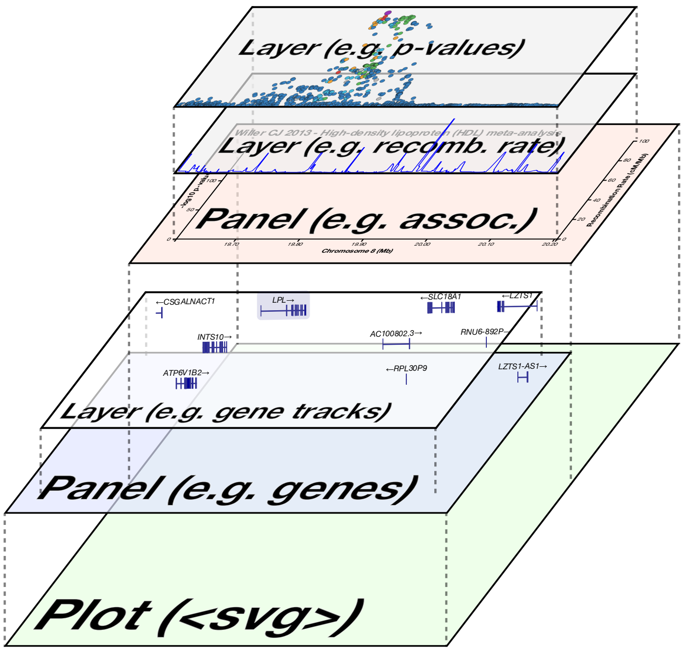


Figure S1: LocusZoom plots are generated from a set of composable rendering elements that can be superimposed (data layers) or stacked in the y-direction (separate panels). This modular architecture allows for customizable and information-dense visualizations.

### Comparison of tools and features

LocusZoom.js incorporates many useful features based on the classic LocusZoom "standalone" command line interface (CLI) (Pruim *et al.*, 2010). For example, the region plot view is quite similar, and there are similar annotation options.

An important difference lies in the underlying design philosophy: the original command line tool was designed to be used on arbitrary data sets stored locally, and therefore assumes that the end user has complete control over every aspect of the visualization. By contrast, LocusZoom.js was designed for data sharing portals or interactive web applications. In this context, it is typically the web application developer (not the end user) who drives many of the display decisions. Both strategies have strengths and weaknesses, and LocusZoom.js aims to leverage the advantages of a web application, rather than exactly duplicating the original command line tool.

In the original LocusZoom CLI, given a sufficiently complex command line string, almost any feature can be activated for a given task; given sufficient modification of the input data, almost any annotation can be added. However, the results are provided as static images, and exploring many regions can be slow and tedious. Updates to the underlying data are not automatically retrieved, which means that visualizations are frequently outdated or missing new annotation information. Emphasis on command line usage and a custom annotation syntax can serve as a barrier to new users. The original LocusZoom.org upload site was built on the command line tool, and thus echoes these limitations. For example, the same file may need to be uploaded multiple times in order to explore a large number of loci, and error messages due to an unrecognized file format can be unintuitive.

With LocusZoom.js, a graphical user interface makes it easy to explore a dataset via sensibly chosen default behaviors. Some features, such as LD reference variant or population, can be modified in real time via user interaction. Individual applications that embed LocusZoom.js as a widget can customize the set of data types and display options to address a particular research question (see Table S1 and Figures S2-S5 for examples). Generally, data have been pre-processed and harmonized prior to viewing, making it easy to explore multiple loci within the same dataset. Web applications are also extremely well suited to sharing results, since collaborators can explore the entire dataset and are not limited to a set of pre-selected loci, as is the case if sharing static PDFs. These advantages encourage collaboration and reduce artificial barriers to entry from command line syntax, but tend to emphasize solving the most common problems at the expense of customization in the most advanced use cases.

In the years since the original LocusZoom CLI and website, other web-based visualization tools have emerged, most often to solve a particular research question within a single web application. Compared to other similar plotting tools (Dadaev *et al.*, 2016; Machiela and Chanock, 2018; Watanabe *et al.*, 2017), the embeddable, client-side rendering approach of LocusZoom.js provides more advanced options for interactive exploration and privacy. For example, when rendering is handled server-side, sensitive data must be shared with a third party that controls that server. LocusZoom.js can provide greater privacy by rendering the data from a local tabix-indexed file, or by embedding in a private intranet site that retrieves data from an API completely under the control of the user. Used as an embeddable library and customizable plugins, LocusZoom.js can expose a wider potential range of annotation options and comparisons than any single purpose-built visualization tool. In both features and privacy options, significant efforts are made to allow incremental adoption based on user needs.

| Feature | LocusZoom.js | LocalZoom | my.locuszoom.org |
|---|---|---|---|
| GWAS region plots | * | * | * |
| Display nearby genes | * | * | * |
| Tooltips link to more information | * | * | * |
| Select LD population | * | * | * |
| Supports GRCh37 and GRCh38 | * | * | * |
| Save as SVG or PNG image | * | * | * |
| Label variants of interest | * | * | * |
| Pan and zoom to change locus | * | * | * |
| Interactive display filters | * | * | * |
| Switch between display modes | * | * | * |
| Compare to GWAS catalog | * | * | * |
| Calculate & display credible sets | (*) | * | * |
| Run rare variant aggregation tests | (*) | | |
| Chromatin co-accessibility | (*) | | |
| BED track intervals | (*) | | |
| Forest plot | (*) | | |
| PheWAS rendering | * | | |
| Embed in any web site | * | | |
| Stack multiple GWAS in one view | * | * | |
| Plot data without uploading | (*) | * | |
| Export data as CSV file | | * | * |
| Compare to a published PheWAS | | * | * |
| Search by gene, rsID, or coordinates | | * | * |
| Batch view mode | | * | * |
| Manhattan plot | | | * |
| Frequency-stratified QQ plot | | | * |
| Summary table of top loci | | | * |
| Identify nearest gene for top loci | | | * |
| Annotate rsID for each variant | | | * |
| Share user-defined datasets | (*) | | * |
| Private links for restricted sharing | | | * |
| Link to an exact region | (*) | | * |
| Explore public studies | * | | * |

Table S1: Comparison of user-facing features for each of the ways to use LocusZoom.js. Symbols: * = yes; (*) = feature implemented as a plugin / extension feature. In this table, three basic tiers are presented: features available in the core visualization library (top), features available in LocalZoom (middle), and features available in the fully managed my.locuszoom.org upload service (bottom).

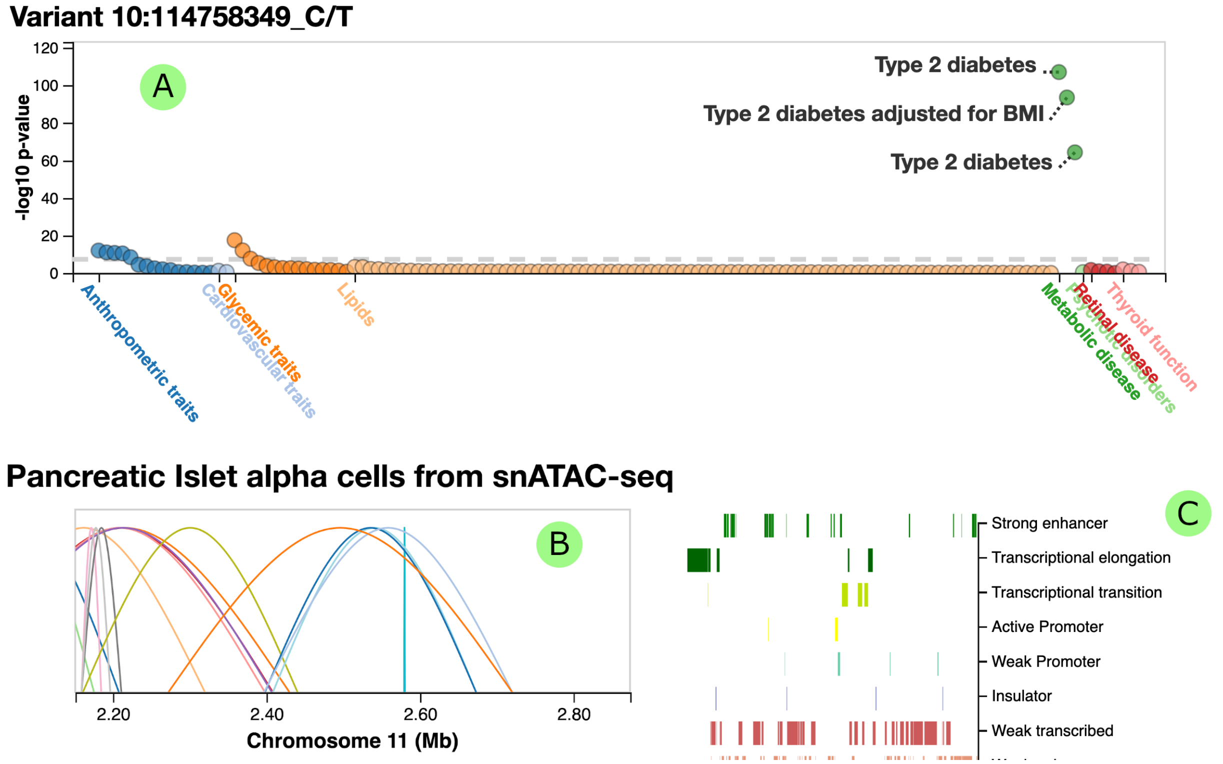


Figure S2: In addition to the familiar regional scatter plot of GWAS summary statistics, LocusZoom supports a wide range of visualization types (see Table S1 for a full list). Examples shown: (a) a single-variant PheWAS scatter plot, (b) chromatin co-accessibility loops, and (c) BED track interval annotations for chromatin state.

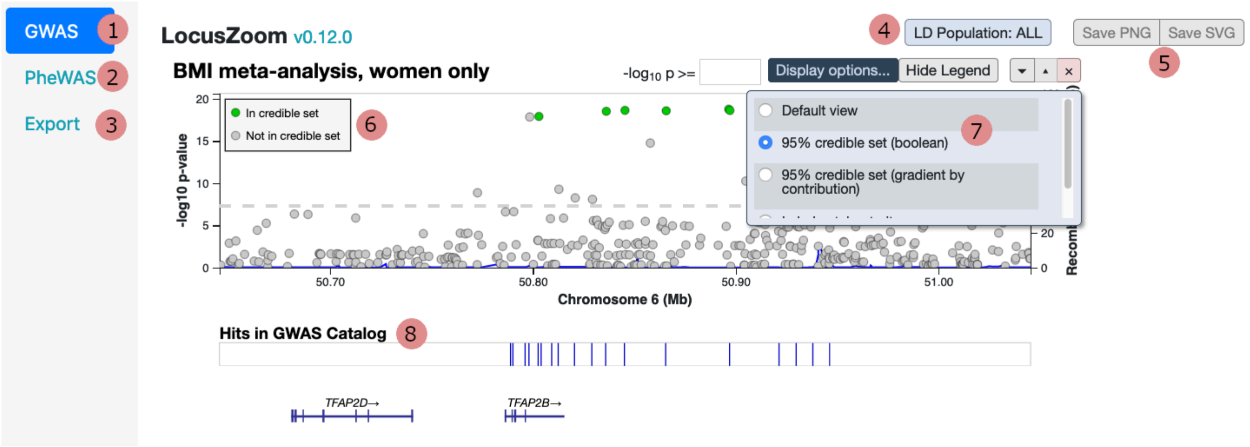


Figure S3: The LocalZoom user interface provides many interactive features for exploring datasets. Separate tabs show data as (1) a classic LocusZoom GWAS region plot, (2) a PheWAS plot to put results in context of a UK BioBank PheWAS, or (3) exporting the summary statistics in this region to a table or CSV file for follow-up analysis. Toolbar buttons allow the user to (4) select one of several linkage disequilibrium reference populations or (5) export the current view as SVG or PNG. (6) A credible set is automatically calculated based on the region in view and (7) the user may interactively change whether or how points are rendered. Additional annotations include (8) a quick view of which genetic variants from this dataset have known significant entries in the GWAS catalog.

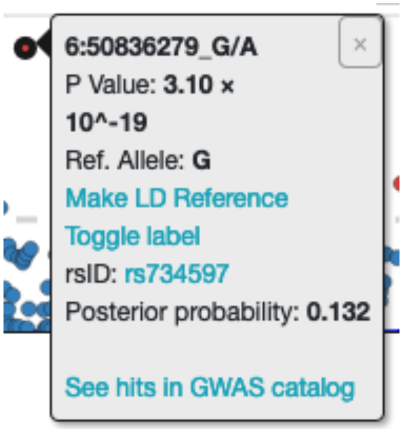


Figure S4: A tooltip appears when the mouse pointer moves over data elements. Tooltips can be used to show additional information (such as rsID), trigger interactive behaviors (change LD reference), or link to outbound resources (GWAS catalog).

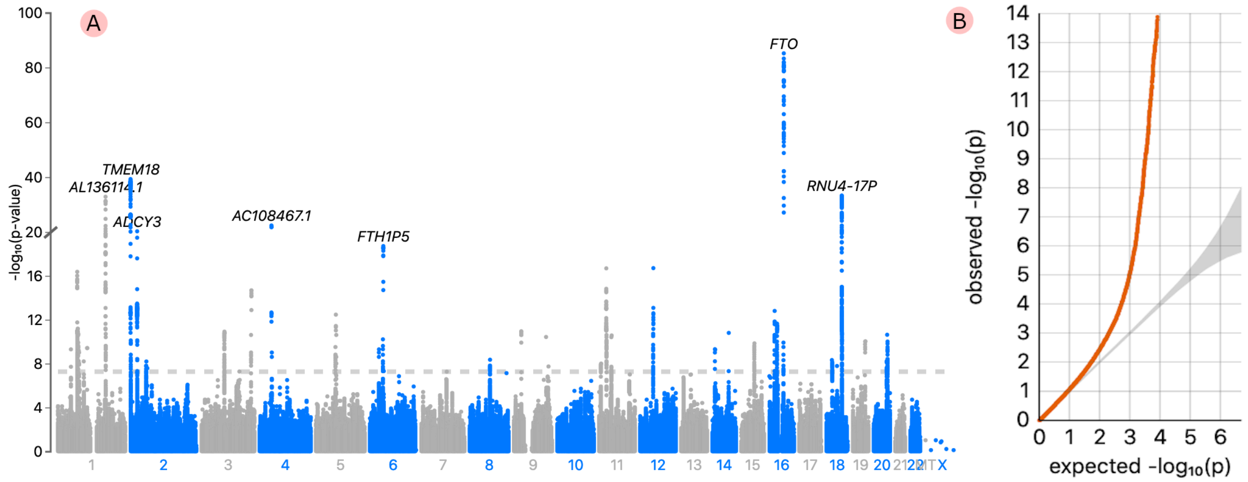


Figure S5: The my.locuszoom.org upload service provides a number of summary and visualization features, such as Manhattan and QQ plots generated from the data. Variants are automatically annotated with rsID and nearest gene. The multiple layers of visualization on this site are connected: clicking on a variant takes the user directly to a LocusZoom region plot view for that signal.
